# Morphological, pathological and phylogenetic analyses identify a diverse group of *Colletotrichum* spp. causing leaf, pod, and flower diseases on the orphan legume African yam bean

**DOI:** 10.1101/2024.04.03.587868

**Authors:** Olaide Mary Ogunsanya, Moruf Ayodele Adebisi, Akinola Rasheed Popoola, Clement Gboyega Afolabi, Olaniyi Oyatomi, Richard Colgan, Andrew Armitage, Elinor Thompson, Michael Abberton, Alejandro Ortega-Beltran

## Abstract

African yam bean (AYB; *Sphenostylis stenocarpa* Hochst. Ex A. Rich.) is an underutilized legume indigenous to Africa. The crop has great potential as it can enhance food security and its chemical composition offers nutritional and medicinal opportunities. However, the low grain yield caused by fungal diseases, including pod blight and leaf tip dieback, deters farmers from large-scale cultivation. The causal agents of pod and leaf tip dieback on AYB are largely uncharacterized. To determine the prevalence of fungal diseases affecting leaves, pods, and flowers of AYB, a survey was conducted in 2018 and 2019 in major AYB-growing areas in Nigeria. Leaf tip dieback, flower bud rot, and pod blight were the most common symptoms. Morphological and molecular assays were conducted to identify the causal agents of the observed diseases. In all the samples examined, fungi from eight genera were isolated from leaves and pods showing disease symptoms. However, Koch’s postulates were fulfilled only for fungi belonging to the *Colletotrichum* genus. Fungi from the other seven genera did not produce disease symptoms in healthy AYB tissues. Several *Colletotrichum* isolates were characterized by sequencing the ITS, glyceraldehyde-3-phosphate dehydrogenase, calmodulin, and ApMAT loci. A combined phylogenetic analysis revealed four *Colletotrichum* species: *C. siamense*, *C. theobromicola,* and *C. fructicola*, which were recovered from the diseased leaves, and *C. truncatum*, which was recovered from diseased pods and buds. Our results are useful to gear efforts to develop integrated management strategies to control diseases affecting AYB in Nigeria and other parts of Africa. The availability of such strategies may stimulate greater cultivation of AYB to contribute to diet diversification, which has been repeatedly advocated by a range of stakeholders to increase food security and the prosperity of smallholder farmers.

## INTRODUCTION

African yam bean (AYB; *Sphenostylis stenocarpa* Hochst. Ex A. Rich.) is a tuberous legume that belongs to the *Fabaceae* family. The crop has the capacity to withstand climatic stresses such as heat and drought, thrives well in marginal soils, and improve soil quality, thereby possessing great potential to enhance food security in various African countries (Nnamani et al. 2017, 2021; Ojuederie and Balogun 2019). However, AYB is underutilized. African yam bean produces two organs of economic importance, grains, and tubers, although not all AYB accessions produce tubers (Adewale and Nnamani 2022). African yam bean is consumed based on regional preferences and beliefs. In West Africa, grains are consumed, but many farmers are unaware that AYB can tuberize, and many of those who are aware believe the tubers are poisonous. In contrast, in East Africa, the tubers are consumed, but many farmers believe the grains are poisonous (Adewale and Nnamani 2022; Nwokolo 1996; Potter and Doyle 1992). However, the grains and tubers of AYB have enormous benefits. They are safe for human and livestock consumption and have nutritional and medicinal benefits (Christopher et al. 2013; Nwankwo et al. 2018; Ojuederie et al. 2020). The tubers are rich in crude protein, total ash, and fat (Konyeme et al. 2020). The grains are rich in minerals, vitamins (Ajibola and Olapade 2016), and fiber (Anya and Ozung 2019). In traditional medicine administered in Enugu state, Nigeria, AYB grains are used to treat insomnia, measles, and diabetes (Nnamani et al. 2021).

Fungal diseases are one of the factors deterring farmers from large-scale cultivation and germplasm regeneration of AYB (Afolabi et al. 2019; Ameh and Okezie 2005). Several diseases of AYB including powdery mildew, leaf spot, stem rust, wilt (Ameh and Okezie 2005), pod blight, flower bud rot, and tip dieback have been reported (Afolabi et al. 2019). In AYB fields managed by researchers of the International Institute of Tropical Agriculture (IITA), located in Southwest Nigeria, tip dieback, wilt, flower bud rot, and pod blights were diseases associated with AYB (O. Ogunsanya, unpublished). These observations raised questions on whether these diseases are common to AYB in all actively AYB-growing regions of Nigeria and what pathogens are responsible for these diseases.

Most reports on fungal diseases associated with AYB have been informal or have not focused on the identification of the causal agents of the diseases. For example, Afolabi et al. (2019) found that fungi from 13 genera were associated with AYB flower bud rot and pod rot diseases, but Koch’s postulates were not conducted. Accurate identification of causal agents of important diseases of AYB is crucial for disease management and breeding purposes. To accurately identify microorganisms associated with a disease, a polyphasic approach is required. Relying on a single identification method may not provide sufficient information to correctly identify the causal agent of a disease (Cai et al. 2009; Simões et al. 2013). Thus, the objectives of the current study were to identify diseases associated with AYB in major AYB-producing areas in Nigeria and to characterize the pathogenic fungi employing a polyphasic approach composed of morphological, phylogenetic, and pathogenicity assays.

## Materials and Methods

### Sample collection

A survey was conducted between 2018 and 2019 in major AYB growing areas in Nigeria to investigate fungal diseases associated with AYB. Samples of diseased AYB were collected from 36 farmers’ fields in Enugu, Ebonyi, and Abia (located in South East Nigeria), and Cross River states (located in South South Nigeria; Fig. 1), as well as from two AYB research fields in IITA-Ibadan in Oyo state (South West Nigeria; Fig. 1). At each survey site, a zigzag transect (area = 3 m^2^) was used to randomly select 10 plants about 5-months old for visual examination and sample collection. The pods, leaves, and flower buds were visually examined for signs of fungal infection: necrotic lesions or discoloration on leaves; browning on flower buds; and rot or lesions in pods. A total of 360 leaf samples (10 per field) were collected from farmers’ fields, placed in appropriately labelled bags, and transferred to a plant press for preservation. Due to lack of resources to keep plant materials fresh and the distance from the laboratory, only leaf samples were collected from farmers’ fields in the Southeast/Southsouth. Information of the specific AYB accessions sampled was not available. Various sample types (20 leaves, 20 diseased pods, and 10 flower buds) were collected from 20 AYB accessions in the research fields in Ibadan because of proximity to the laboratory and access to ice to preserve the samples.

**Figure 1.**
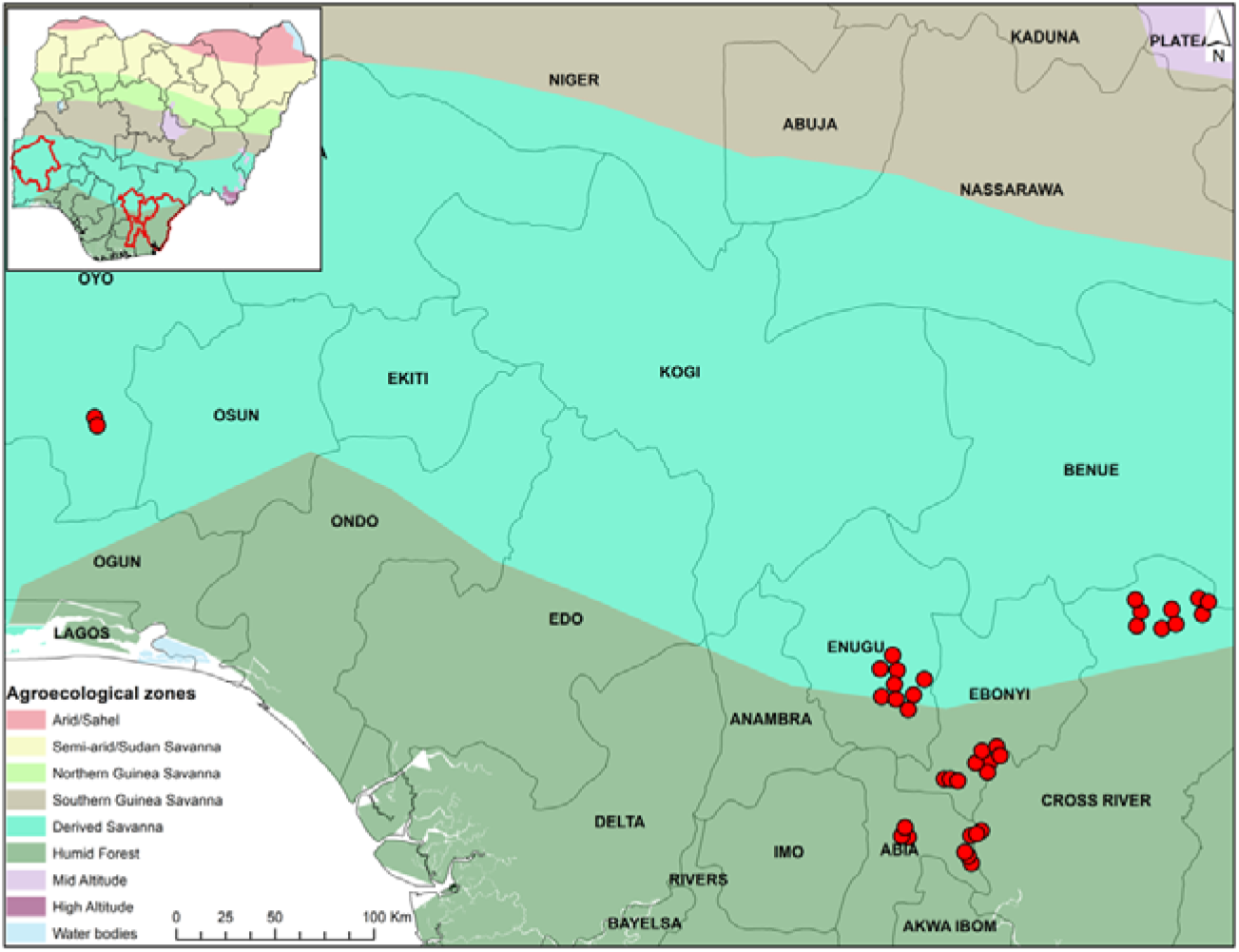
Area of study in Nigeria, with red circles representing the locations of African yam bean fields sampled between 2018 and 2019 with the aim of identifying diseases prevalent in these regions and elucidating their respective causal agents.

### Fungal isolation

Fungal isolates were recovered from infected AYB tissues (flower buds, pods, and leaves). Appropriate sections (∼6 mm^2^ piece comprising 1/3 of diseased and 2/3 of the healthy tissue) were excised with a sterile scalpel and surface sterilized in 50% (v/v) NaOCl for 30 s. Samples were then triple rinsed with sterile distilled water, and then blotted dry using sterile paper towels in a Class II microbiological safety cabinet. With the aid of a mounting needle, four segments of the sterilized plant tissues were placed in Petri dishes containing acidified potato dextrose agar (PDA + lactic acid 0.1%). Petri dishes were incubated for 3 days at room temperature (25 - 28°C). A maximum of four discrete colonies of fungi per sample were sub-cultured onto PDA and incubated at 25 - 28°C for 5 days. Then, cultures of each fungal isolate recovered were single-spored using the method of Goh (1999), with minor modifications. The procedure entailed transferring spore masses with the aid of sterile toothpick and suspending them into sterile distilled water. The suspensions were then diluted to 4 × 10^6^ spores ml^-1^. Subsequently, 80 µl of each diluted suspension were spread evenly onto the surface of water agar plates which were incubated overnight at 25°C.

Under the dissecting microscope (Wild Heerbrugg, Switzerland), single germinated spores were identified. Each individual spore, along with its surrounding agar block, was carefully excised using a sterile scalpel, and carefully transferred onto new PDA plates. Plates were incubated at 25°C to allow for fungal growth and sporulation.

Axenic cultures were identified through their morphology on PDA after 8 days of incubation and then conidia were viewed using a compound microscope (Olympus, BX51) at 40× magnification. The pure cultures of each isolate were maintained in the refrigerator at 4°C on Petri dishes and slopes containing PDA for about 14 days. Separate cultures were made for saving in 20% glycerol and stored at −80°C for long term preservation.

### Planting material for pathogenicity assays

One accession of AYB, TSs 1, was used for the experiments. TSs 1 is frequently used in diverse experiments in IITA and therefore selected for the current study. Seeds of TSs 1 were obtained from the Genetic Resources Centre (GRC) of IITA. Two seeds were sown in 8 kg-pots filled with sterilized field soil; no fertilizer was added. The plants were grown in a screenhouse at IITA. Two months after planting, healthy leaves were carefully excised, placed in transparent bags, and sterile distilled water was slightly sprinkled on the leaves in the bags for immediate transportation to the laboratory.

### Box preparation for detached leaf assay (DLA)

Clear plastic boxes (23 × 31 × 10 cm; Pioneer Plastics, USA) with lids were sterilized in 50% NaOCl for 5 min. Then, the boxes were rinsed twice with sterile distilled water. Immediately the boxes were transferred into a Class II microbiological safety cabinet. The boxes were left to properly drain, after which the ultraviolet light in the biosafety cabinet was turned on for 10 min for further decontamination. Sterilized paper towels and cotton wools were laid in the base of the sterilized boxes. Following this, the paper towels and cotton wools were soaked with 100 ml sterilized distilled water amended with 200 µl Hexacal (0.02% fungicide; Farmpays, Nigeria) to inhibit growth of saprophytic fungi. Detached leaves remained viable for up to 16 days.

### Inoculum preparation

Fungal isolates recovered from diseased AYB tissues (leaves and pods) were grouped based on their cultural characteristics. Twenty-five representative isolates recovered from leaves and six representative fungal isolates recovered from pods were selected to be included in DLA. Conidia of each of the fungal isolates tested in DLA were washed off from 7-day-old PDA cultures by adding 2 ml sterile distilled water amended with TWEEN®20 (0.01% v/v). These suspensions were transferred to sterile 10 ml vials. The concentrations of filtrates were adjusted to 1×10^6^ conidia ml^-1^ in sterile distilled water using a haemocytometer (Improved Neubauer Bright Line; Hausser Scientific) and a compound microscope (Leitz Wetzlar, Germany; Laborlux S, magnification 40×).

### African yam bean leaf surface sterilization, inoculation, and disease assessment

For the detached leaf assay (DLA), leaves were submerged for 30 s in distilled water amended with 100 ppm/l of Vertimec (acaricide; Syngenta®) to prevent mite infestation in the laboratory. Leaves were then surface sterilized with 1% NaOCl and dried, as described above in the fungal isolation section. Thereafter, four leaves were carefully placed in a labelled sterile clear plastic box with the adaxial side placed on wet paper towel while the abaxial side was inoculated using a pipette tip without wounding the leaves (Bankole et al. 2022). For each of the 25 evaluated isolates, 10 µl of the 1×10^6^ conidia ml^-1^ suspension was inoculated onto the top and bottom regions of the leaf abaxial surface as described by Bankole et al. (2022). The experiment included mock-inoculated controls using TWEEN^®^20 (0.01% v/v). The boxes were sealed with plastic wrap cling film to help keep the chamber humid and incubated at room temperature (∼25°C) on the benchtop with a cycle of 12 h of light and 12 h of dark for 8 days. Inoculated AYB leaves were examined every 2 days for disease progression. The disease was scored under a sterile microbiological safety cabinet, with containers remaining sealed to avoid contamination. Necrotic lesions on leaves were visually assessed by the percentage area of leaf infected and were scored on a scale of 1-5 where 1 = no disease symptom, 2 = up to 10 %, 3 = 10 – 25 %, 4 = 26 – 50 %, 5 = > 50 % of leaf surface area showing symptoms. This scale was adapted from the one reported by Nwadili et al. (2017). At the end of the incubation period (8 days after inoculation), the inoculated isolates were re-isolated from the symptomatic AYB tissues and transferred to PDA to confirm their identity as the causal agent of the disease. The experiments were set up in a Completely Randomized Design (CRD) with four replications per isolate. The DLA was conducted twice.

### African yam bean pod collection, laboratory inoculation and disease assessment

In this experiment, 50 AYB seeds of TSs 1 were sown in a research field at IITA. Plots were formed by single 4-meter-long ridges spaced 0.75 m apart. The seeds, treated with mancozeb 80% wettable powder (WP), were planted 0.5 m apart on the ridges. Three weeks after the emergence of seedlings, triple superphosphate fertilizer was applied at a rate of 50 kg ha^-1^. Every two weeks, a mixture of Cypermethrin 30 g/l + Dimethoate 250 g/l EC and Mancozeb 80% WP was applied at a rate of 200 ml and 200 g/20 l water, respectively. Standard agronomic practices such as weeding (when necessary), and staking (after three weeks after planting) were conducted at the appropriate time.

Healthy AYB pods (5-month-old) were carefully excised and placed in a transparent bag which was lightly sprinkled with water before transporting to the laboratory. In the laboratory, the detached pods were sterilized for 1 min in distilled water amended with a drop of Vertimec, rinsed with sterile distilled water and blotted dry. Thereafter, three pods were carefully placed in each labelled sterile clear plastic boxes. Conidial suspensions for each of the six isolates recovered from pods, belonging to representative groups, were prepared as above, and 10 µl of 1×10^6^ conidia ml^-1^ suspension was inoculated, without wounding, using a pipette tip onto three different points of a healthy pod (top, middle, and bottom regions of the pod). The experiment included mock-inoculated controls using sterile TWEEN^®^20 (0.01% v/v), which was prepared by adding 0.1 ml of TWEEN^®^20 to 1 l distilled water and then sterilized. The boxes were sealed with plastic cling wrap film to help keep the chamber humid and incubated at room temperature (∼25°C) on the benchtop with a photoperiod cycle as above for nine days. Inoculated pods were examined every third day for disease progression.

The disease was scored under a sterile microbiological safety cabinet, with containers remaining sealed until the end of the experiment to avoid contamination. Necrotic lesions on pods were assessed by the percentage area of pod infected and were scored on a scale of 1-5: 1 = no disease symptom, 2 = 1 – 10 %, 3 = 11 – 20 %, 4 = 21– 50 %, 5 = >50 % of pod area showing symptoms (adapted from Nwadili et al. 2017). At the end of the trial, the inoculated isolates were re-isolated from infected AYB pods and transferred onto PDA to confirm their identity as the causal agent of the disease. The experiments were set up in a CRD with four replications per isolate. The detached pod assay was conducted twice.

### Extended characterization of pathogenic fungi

Isolates that were pathogenic in the detached leaf and pod assays, all showing characteristics of the *Colletotrichum* genus, were subjected to additional morphological characterization. Mycelia plugs were aseptically collected from actively growing edges of 7-day-old single conidium sub-cultures using sterile 5 mm diameter plastic cork borers and placed individually at the center of PDA plates. The cultures were incubated at 25 - 28°C, with three replicates per isolate. The colony growth rate was recorded every 2 days for 8 days. The conidia and appressorium shape were examined using a compound microscope (Olympus, BX51). Appressoria were produced using the slide culture technique (Cai et al. 2009). Conidial size (length and breadth) was measured using a compound microscope with a graduated eyepiece (Leitz Wetzlar, Laborlux S, Germany).

### Statistical analysis

The disease score values of *Colletotrichum* isolates evaluated in pathogenicity assays were subjected to Kruskal-Wallis test and Dunn-Bonferroni tests (post-hoc test) was used for the pairwise multiple comparisons using GraphPad Prism v7.0 (La Jolla, CA, USA). Student’s *t*-tests were used to compare the growth rate and spore size of *Colletotrichum* isolates. Graphs depicting the mean values with corresponding standard errors of the mean (± SEM) were produced using GraphPad Prism v7.0.

### Genomic DNA extraction

Mycelia plugs (∼3 mm; 1 per isolate) of five-day old monosporic cultures of fungal isolates found to be pathogenic were independently inoculated in potato dextrose broth (Difco^TM^, France) and shaken at 300 RPM for 5 days using a dual-action KL 2 orbital shaker (Edmund Buhler 7400, Tubingen). Mycelia were harvested from broth using vacuum filtration. The setup was as follows; a sterile porcelain Buchner funnel was placed on top of a sterile conical filtering flask to collect the filtrate. A vacuum pump was connected to the filter flask with appropriate tubing. Sterile filter paper of appropriate size was cut to fit the funnel. The mycelia was gently poured into the filter funnel and filtered until most of the liquid passed through. Thereafter, the mycelia were rinsed with sterile distilled water to remove residual medium. Then, a sterile spatula was used to collect the mycelia into a sterile falcon tube and stored at 20°C. Two different methods were used for DNA extraction: Zymo Research-Quick DNA Fungal/Bacterial Miniprep Kit following the manufacturer’s recommendation and Shorty buffer method (Harrison and Thompson, 2020; Edwards *et al.,* 1991).

The shorty buffer method is as follows: About 40 mg of mycelia were collected into Eppendorf tubes and subsequently pulverized in liquid nitrogen using sterile micro pestles. Following this, 500 μl sterile Shorty buffer (200 mM Tris HCl pH 8; 400 mM LiCl; 25 mM EDTA pH 8; 1% w/v SDS) were added to the powdered mycelia. The mixture was vortexed for 10 sec and then centrifuged at 16,000 g for 5 min. In separate Eppendorf tubes, 350 μl isopropanol were carefully added. Then, 350 μl of the supernatant obtained from the centrifugation step was pipetted into the isopropanol-containing tubes. The mixture was gently inverted for 30 sec to allow DNA precipitation and then centrifuged at 16,000 g for 15 min to effectively pelletize the DNA. The supernatant was removed without disturbing the DNA pellet, and 500 μl 70% ethanol were added for DNA washing. A subsequent centrifugation step at 10,000 g for 2 min facilitated the separation of the ethanol, after which the supernatant was carefully pipetted off. To ensure the removal of residual isopropanol and ethanol, tubes were inverted and incubated at 37°C for 15 min. Finally, the DNA pellets were resuspended in 30 μl sterile TE buffer (0.5 ml pH 8; 0.1 ML EDTA; 50 ml sterile distilled water) to complete the extraction process.

### Polymerase Chain Reaction (PCR) and gel electrophoresis, DNA purification, and sequencing

The primers used in the current study targeted the Internal Transcribed Spacer (ITS), glyceraldehyde-3-phosphate dehydrogenase (GAPDH), calmodulin (CAL), and Apn2-MAT 1-2 intergenic spacer (ApMAT) regions of the examined isolates. These primers have proven effective in identifying *Colletotrichum* species (Hassan et al. 2018) and therefore selected. The primer pairs are listed in Table 1. For all primer pairs, the PCR cocktail mix (25 µl) consisted of 12.5 µl OneTaq® Quick-Load® (2x Master mix; New England Biolabs), 1.5 µl each of forward (10 µM), and reverse (10 µM) primers, 1 µl DNA template, and 8.5 µl Nuclease free water.

**Table 1.**
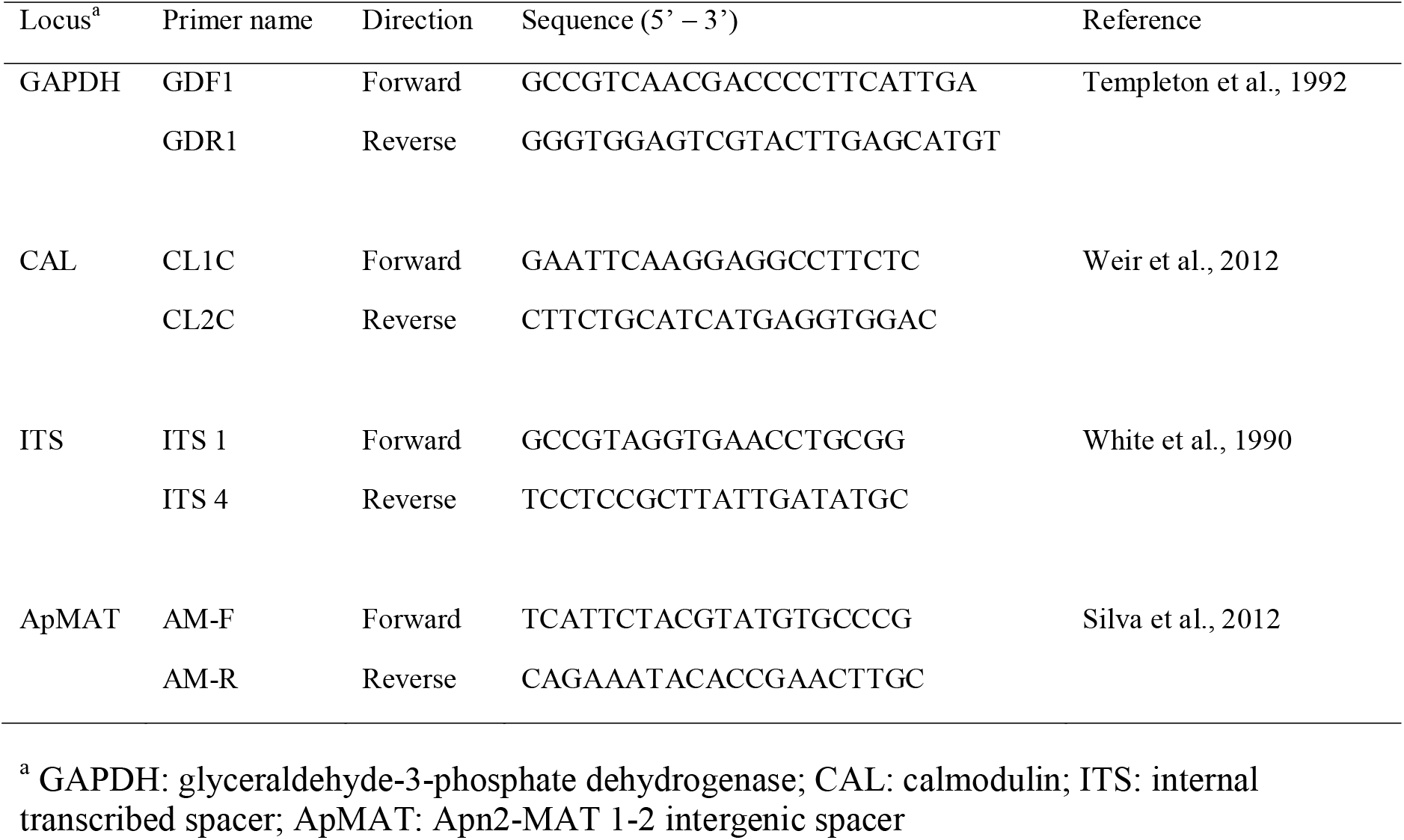
Primers used for the multiple loci analysis of Colletotrichum isolates identified in the current study.

Initially, 42 isolates were subjected to PCR followed by sequencing of the ITS locus to select isolates for the multi-locus sequence analysis. The PCR conditions for ITS, involved initial denaturation at 94°C for 3 min, followed by 30 cycles of denaturation at 94°C for 45 s, annealing at 54°C for 30 s, elongation at 72°C for 45 s, a final elongation step at 72°C for 7 min, and hold temperature at 4°C. PCR products were visualized on 1% agarose electrophoresis gels at 100V for 1 h in 1x Tris-acetate-EDTA buffer. Amplicons were purified using a Monarch Genomic DNA purification kit following the manufacturer’s recommendation, and sequenced bidirectionally by either IITA Bioscience Center (Nigeria), Source Bioscience (United Kingdom), or Eurofins Genomics (Germany).

Based upon the ITS phylogenetic analysis, 13 isolates were further selected for analysis through sequencing portions of GAPDH and CAL loci. The ApMAT locus was examined for the 10 isolates in the *Colletotrichum gloeosporioides* complex (*Cg* species complex) but not for the *Colletotrichum truncatum* isolates due to its specific design for *Cg* species complex (Silva et al. 2012). The PCR conditions remained consistent with those mentioned earlier, except for the annealing temperatures: GAPDH at 55°C, CAL at 59°C, and ApMAT at 62°C.

### Phylogenetic analysis

The raw nucleotide dataset generated from sequencing the ITS locus for each isolate were reviewed, edited, and assembled into consensus sequences using BioEdit Sequence Alignment Editor v7.2.5e (Hall 1999). The ITS sequences of type *Colletotrichum* species were retrieved from NCBI for use as reference. MEGAX v10.2.6 (Kumar et al. 2018) was used to perform multiple sequence alignments and the statistical selection of best-fit models. The nucleotide substitution model with the lowest Bayesian Information Criterion score was considered to describe the substitution pattern the best and used for subsequent phylogenetic analysis. A maximum likelihood phylogenetic trees was constructed using MEGAX v10.2.6 and the analysis was performed with 1000 bootstrap replicates to assess support for the resulting phylogenetic clades. The phylogenetic tree was exported and visualised using Figtree v1.4.4 (Rambaut 2018).

The raw nucleotide dataset generated from sequencing the CAL and GAPDH regions of the 13 selected isolates were subjected to sequence editing and assembly to generate consensus as above. From NCBI, the CAL and GAPDH sequences of the type *Colletotrichum* isolates used above were also retrieved. The nucleotide dataset generated from sequencing the ApMAT region of the 10 *Cg* species complex isolates were subjected to post sequencing analysis as above and the ApMAT region sequences of the same type *Colletotrichum* isolates were retrieved from NCBI (Suppl. Table 1).

The consensus sequences of the 13 isolates were aligned and concatenated using Geneious Prime v2022.2.2 (Biomatters Ltd. 2021), to generate a composite consensus sequence for each isolate that contained the ITS region as well as the CAL, and GAPDH loci, in addition to ApMAT for the 10 *Cg* species complex isolates. A maximum likelihood phylogenetic tree was constructed and exported as above.

## Results

### Field observations

A total of 380 plants were examined across locations. The most common symptoms were leaf tip dieback, flower bud rot, and pod blight (Fig. 2). The frequency of these symptoms varied among locations. However, pods were most likely to show blight (mean = 53%, N = 380) whereas leaf tip dieback (mean = 38%, N = 380) and bud rot (mean = 37%, N = 380) were noted in fewer of the examined samples. The characteristic symptom of leaf tip dieback was a necrotic lesion at the leaf margin which progressed inwardly. A pod blight disease was characterized by irregular-shaped necrotic lesions with distinguishing signs of acervuli (ringed black mass of fruiting bodies) on pods. In addition, some infected pods were twisted, some did not contain any seeds, and others had few seeds. All diseased flower buds were completely brown and non-viable.

**Figure 2.**
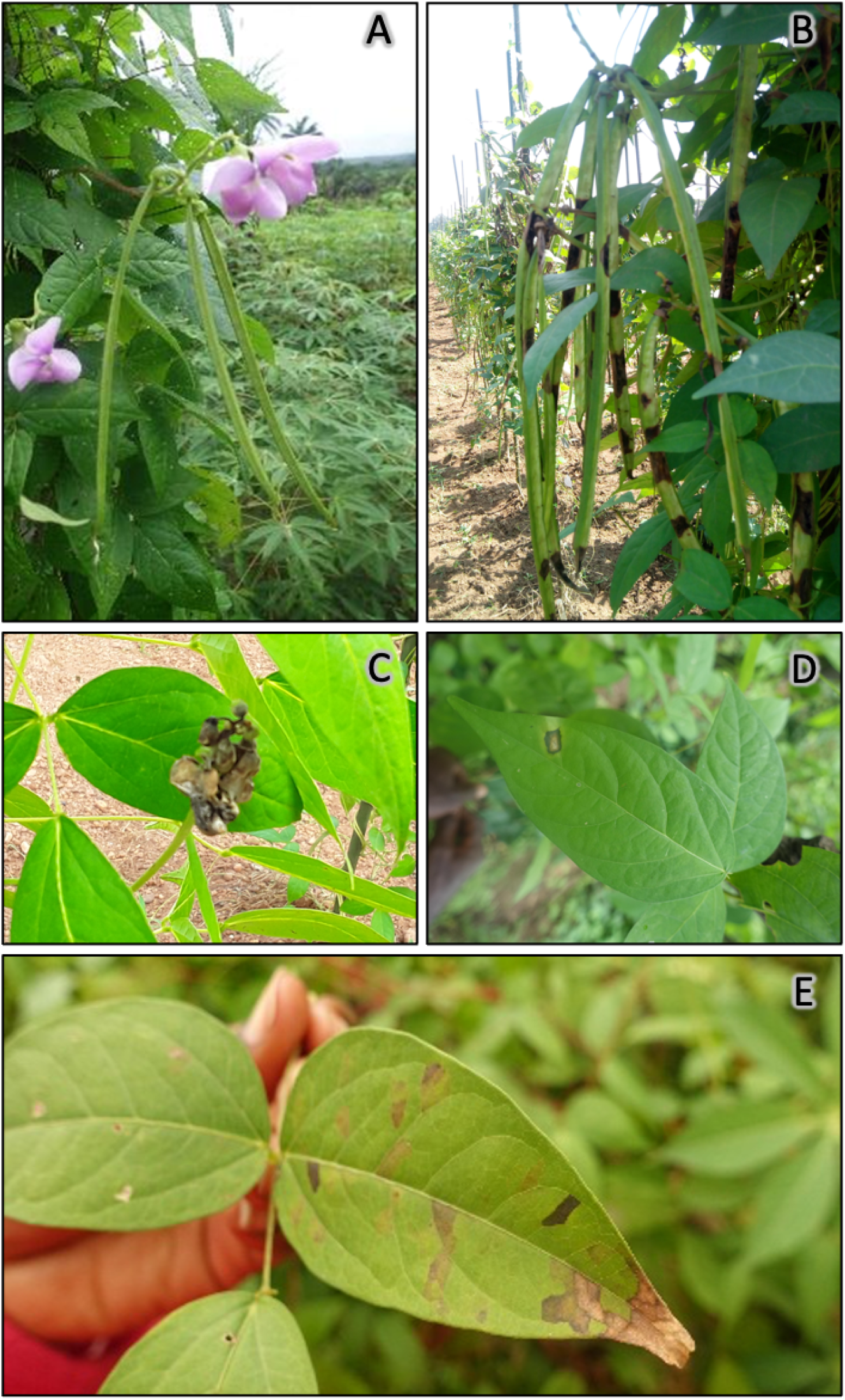
African yam bean plants in the visited fields. A: healthy pods and flowers; B: blighted pods; C: flower showing bud rot; D: leaf blight; E: leaf tip dieback.

### Fungal isolation from AYB tissues

A total of 231 fungal isolates were recovered from infected AYB tissues: 15 from diseased flower buds, 62 from diseased pods, and 154 from diseased leaves. Based on morphological characteristics, eight fungal genera were identified among the recovered isolates. The most frequently isolated fungi from symptomatic leaves were isolates of the *Cg* species complex (44%; Fig. 3). *Fusarium* spp., *Curvularia* spp., and *Pestalotia* spp. were also recovered from infected leaves at frequencies ranging from 10% to 16%. *C. truncatum* composed about 98% and 87% of the fungi recovered from the pods and flower buds (Fig. 3), respectively, but was never recovered from leaves. *Cg* species complex isolates composed 2% of the fungi recovered from the pods (Fig. 3). Other fungal genera occurring at frequencies equal to or less than 6% include *Botryodiplodia* (6%), *Seridium* (5%), *Exserohilum* (2%), and *Drechslera* (1%).

**Figure 3.**
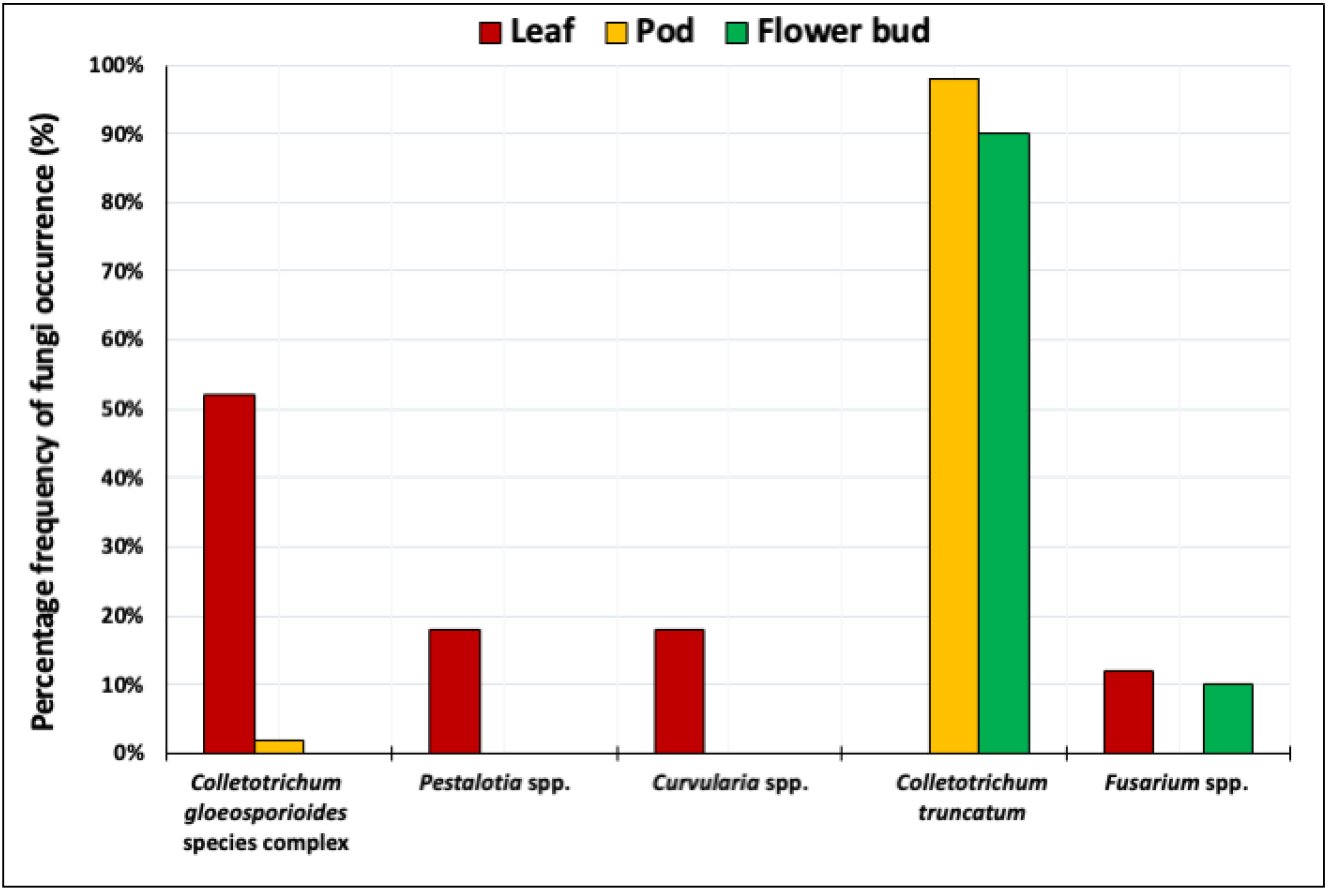
Fungi associated with African yam bean leaves, pods, and flower buds in the sampled areas.

### Pathogenicity assays on detached leaves and pods

All the evaluated isolates belonging to *Cg* species complex (origin information provided in Suppl. Table 2) and *C. truncatum* (all from the research field in Ibadan) caused disease symptoms on the inoculated leaves and pods, respectively, in the laboratory assays. None of the isolates belonging to other genera produced disease symptoms during the 8 days of incubation and therefore did not satisfy Koch’s postulates. The disease scores for all isolates that produced symptoms were significantly (*P* < 0.0001; Suppl. Table 3) different from the symptomless uninoculated control leaves (Fig. 4) or pods (Fig. 6).

**Figure 4.**
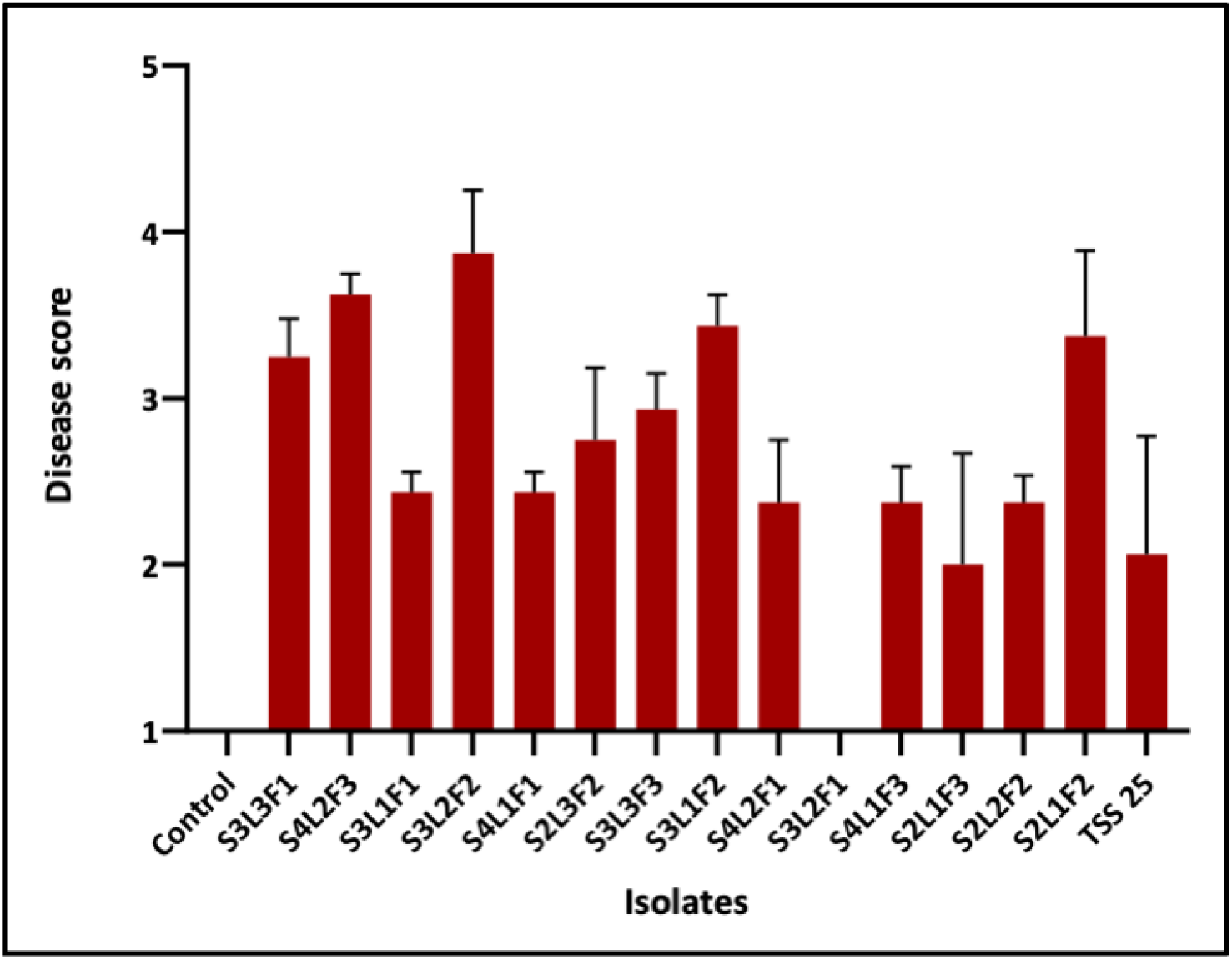
Disease severity scores of 14 isolates belonging to the *Colletotrichum gloeosporioides* species complex when inoculated on African yam bean leaves. The origin of the isolates is provided in Supplementary Table 2.

In the DLA, all *Cg* species complex isolates produced a characteristic progressive irregular black lesion on leaves (Fig. 5). However, there were significant differences in pathogenicity among the 15 *Colletotrichum* isolates (Kruskal-Wallis test, H = 47.18, *P* < 0.0001; Fig. 4). Further, the post-hoc test revealed that while majority of the isolates differ in pathogenicity, five isolates (S3L1F1, S4L2F1, S3L2F1, S4L1F3 and S2L2F2) were not significantly different (*P*-value range = 0.0995 to 0.9999) from the symptomless, uninoculated control leaves. (Fig. 4). Isolate S3L2F1 did not produce disease symptoms on unwounded leaves (Fig. 5) but it was able to cause disease when inoculated on wounded leaves (data not shown).

**Figure 5.**
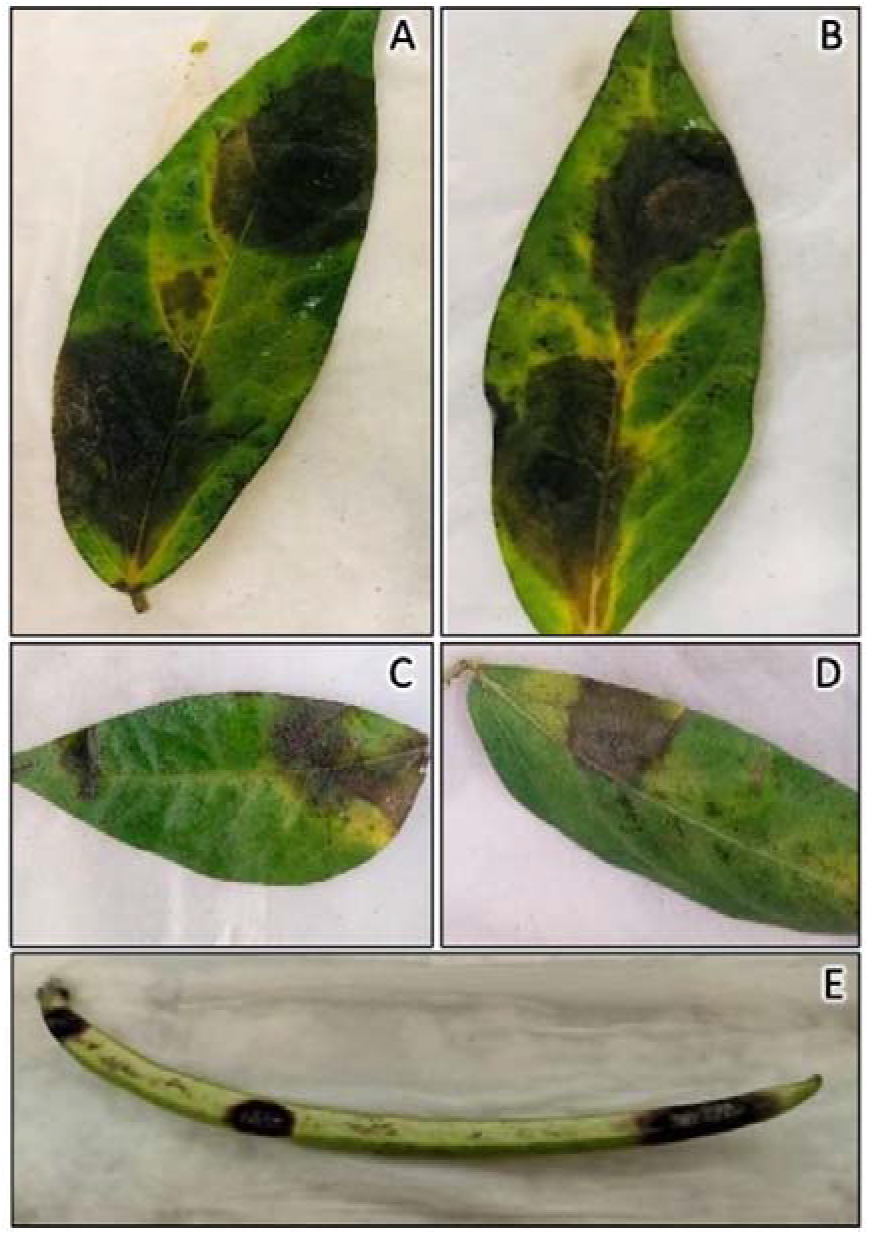
Disease reaction of *Colletotrichum* species inoculated on detached African yam bean leaves and pods. A – D: leaf lesions caused by isolates from the *C. gloeosporioides* species complex (S2L1F2, S2L1F3, S2L3F2, and S4L1F1, respectively); E: pod lesions caused by *C. truncatum* isolate Tss29.

Some isolates induced disease symptoms at 2 days post inoculation (dpi) while some at 6 dpi (Suppl. Table 3). However, at 8 dpi all AYB leaves inoculated with all *Cg* species complex isolates (except S3L2F1) showed anthracnose symptoms. The *C. truncatum* isolates were in general less virulent on AYB leaves (post-hoc test, *P* < 0.001) than *Cg* species complex isolates (Fig. 6). Indeed, the symptoms caused by *C. truncatum* (except TSs 421) were generally minor and not significantly different from the non-inoculated control leaves (Suppl. Table 3).

**Figure 6.**
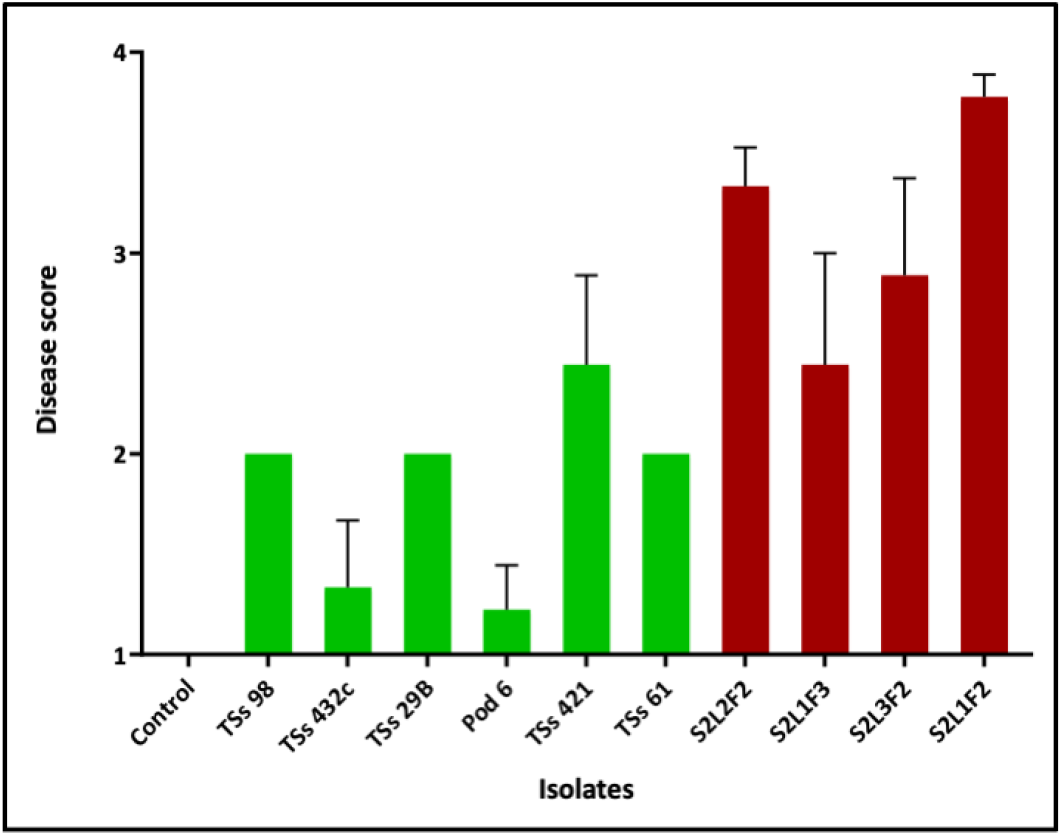
Variation in pathogenicity among *Colletotrichum truncatum* (green bars, all recovered from pods) and *Cg* species complex isolates (red bars, all recovered from leaves) inoculated on African yam bean leaves. Isolates of *C. truncatum* were less pathogenic on leaves.

*C. truncatum* was the sole fungal species included in the detached pod assay because it was the predominant fungus found in pods (Fig. 3) and because there were limited pods for the experiment. On pods, at 3 dpi, all *C. truncatum* isolates produced disease symptoms (Fig. 5). Isolate TSs 432 had the least disease score on AYB pods. The non-inoculated control pods remained symptomless.

### Extended characterization of *Colletotrichum* species

The encountered *Colletotrichum* species were grouped based on conidia shape, rod-shaped in Group A and falcate shaped in Group B (Suppl. Fig. 1). Group A were all *Cg* species complex, predominantly isolated from the leaves while Group B were all *C. truncatum* isolated from pods and flower buds.

There were varying conidia sizes with isolates in Group B having significantly longer (unpaired *t-*test, *P* < 0.0001; *t* = 14.80) conidia than those in Group A. The conidia size of Group B isolates ranged from 7.5 – 22.5 µm × 2.5 – 7.5 µm, whereas in Group A isolates it ranged from 5.0 – 17.5 µm × 2.5 – 5.0 µm (Suppl. Fig. 2). In addition, Group A grew faster than Group B isolates with mycelial growth rate for Group A isolates ranging from 31 to 72 mm at 8 dpi compared to 25 to 57 mm for Group B isolates, but these differences were not statistically significant (*P* = 0.187; *t* = 1.334). The colony morphology of Group A (Fig. 7) and Group B (Fig. 8) isolates on PDA was inconsistent, with some isolates within the same group exhibiting two distinct cultural types on PDA (Suppl. Table 4).

**Figure 7.**
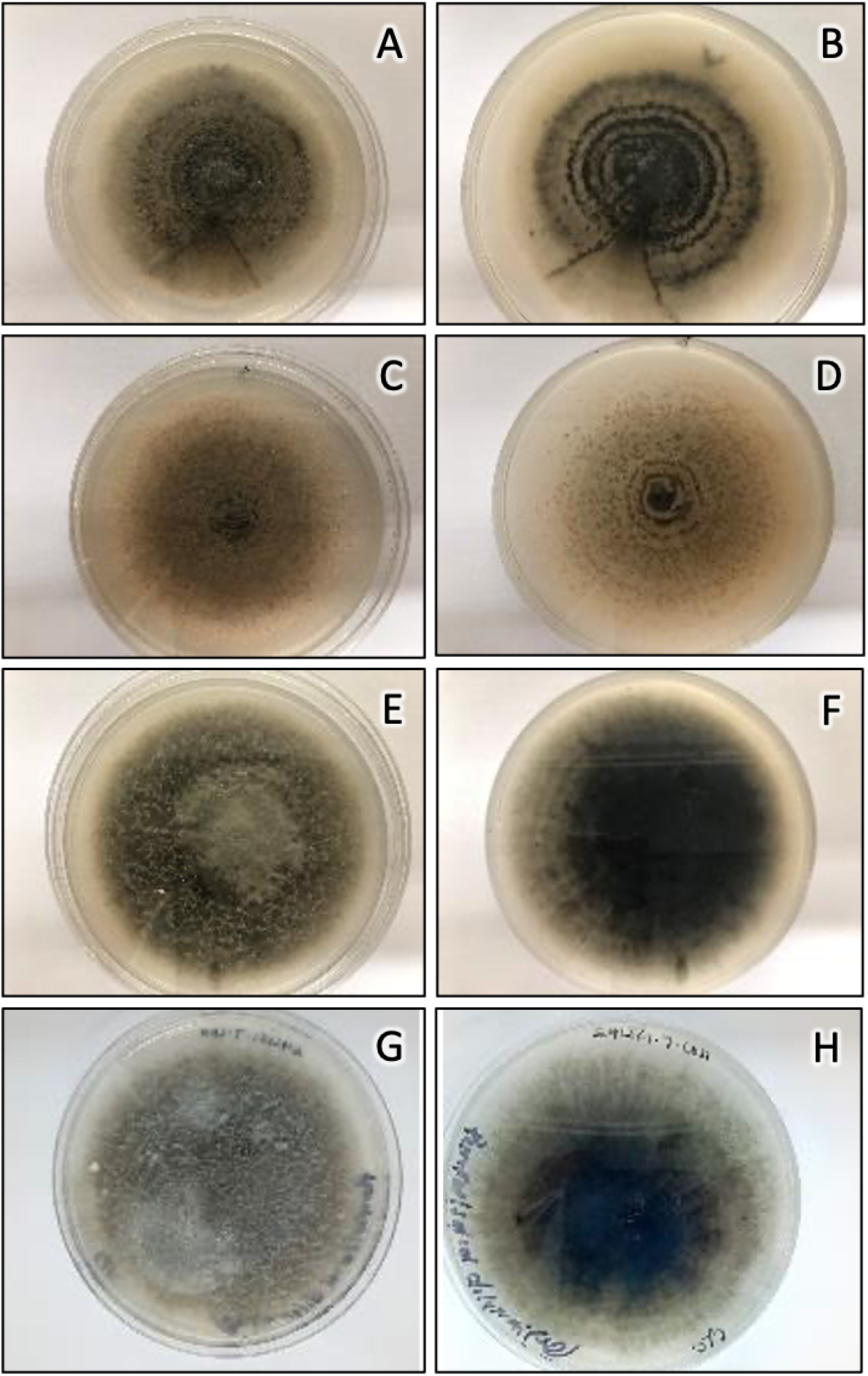
Colony morphology of *Colletotrichum gloeosporioides* species complex isolates on PDA eight days after inoculation. Front and reverse pictures are shown for isolates S2L1F2 (A and B); S2L1F3 (C and D); S2L3F2 (E and F); and S4L1F1 (G and F). All four isolates were obtained from leaves.

**Figure 8.**
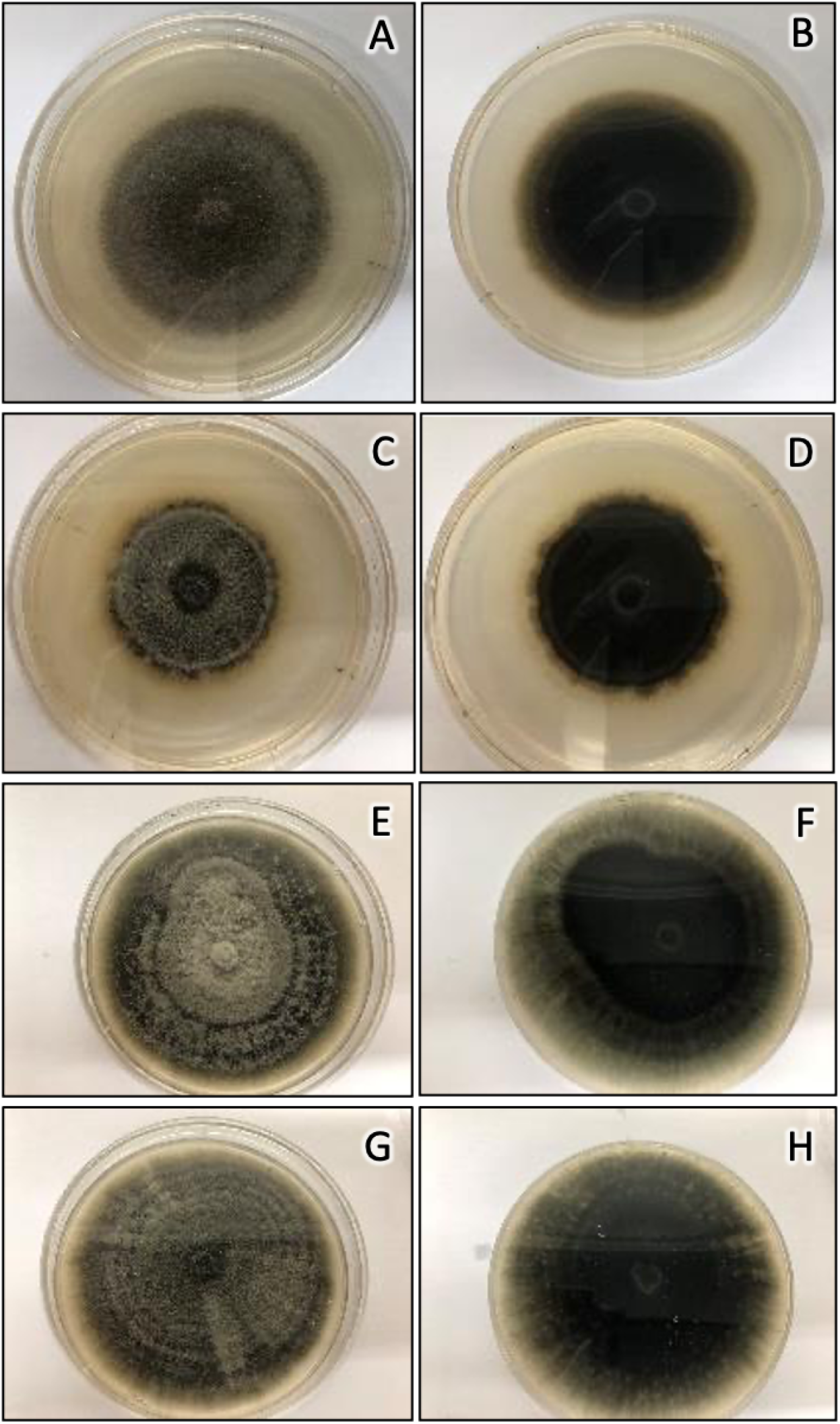
Colony morphology of *Colletotrichum truncatum* isolates on PDA eight days after inoculation. Front and reverse pictures are shown for isolates TSs 29 (A and B); TSs 98 (C and D); Pod 6 (E and F); and TSs 424 (G and F). All isolates four were obtained from pods.

Minor differences were observed among the groups in terms of appressoria shape and color. In both groups, the appressoria were brown to dark brown in colour and lobed to irregular in shape. In addition, only Group B isolates possessed setae. The setae were commonly septate, and cylindrical with sharp tips (Suppl. Fig. 3).

### Phylogenetic analysis

The phylogenetic analysis revealed three species among the *Colletotrichum* isolates causing leaf diseases while members of a single *Colletotrichum* species produced the pod disease. Initially, the ITS locus was amplified and sequenced for 42 isolates with the phylogenetic analysis revealing two clades. However, the ITS phylogeny, while effective in distinguishing the 17 *C. truncatum* isolates from the 25 isolates within the *Cg* species complex clade, provided inadequate resolution to differentiate between species within the *Cg* species complex.

A multi-locus analysis (ITS-GAPDH-CAL) resolved 13 selected *Cg* isolates into four clades (Fig. 9). Isolates S3L2F1, S2L2F2, S3L2F2, S2L1F2, S2L1F3, TSs 57, S2L3F2, TSs 25 clustered in the *C. siamense* group. Isolates S3L3F3 and S4L1F1 clustered in the *C. fructicola* and *C. theobromicola* group, respectively. The *C. truncatum* isolates TSs 29, TSs 432, and Pod6 continued grouping in the same clade, as in the ITS phylogeny tree (data not shown). However, in the four loci analysed (ITS-GAPDH-CAL-ApMAT), S2L3F2 grouped in the *C. fructicola* clade (Fig. 10). Furthermore, within the *C. siamense* clade, S2L1F2, S2L2F2, S2L1F3 (from different Local Government Areas in Cross River) and TSs 57 (from Oyo) formed subclade 1 while S3L2F1 and S3L2F2 (from the same Local Governments Area in Ebonyi) formed subclade 2.

**Figure 9.**
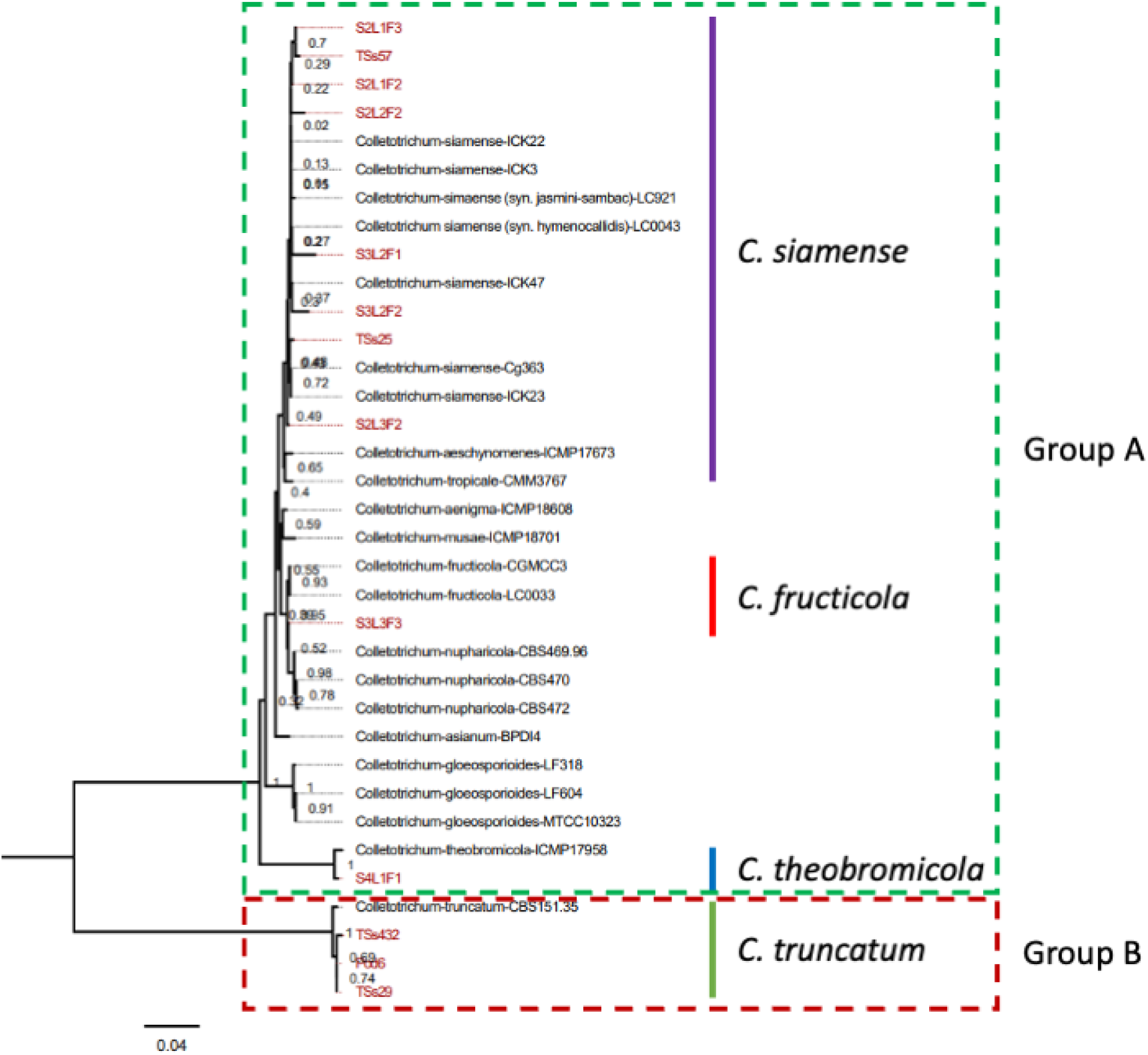
*Colletotrichum* species phylogeny inferred by maximum parsimony from concatenated sequences of the ITS, GAPDH, and CAL loci. The 13 isolates from the current study are highlighted in red. Labels in black are for isolates for which sequences were retrieved from GenBank. The image represents the most parsimonious tree. The bootstraps support values are shown at the nodes. The *Colletotrichum gloeosporioides* species complex isolates, Group A, are located within the dashed box in green while those belonging to *C. truncatum*, Group B, are in the dashed box in red. The tree is rooted with *C. higginsianum* as the outgroup.

**Figure 10.**
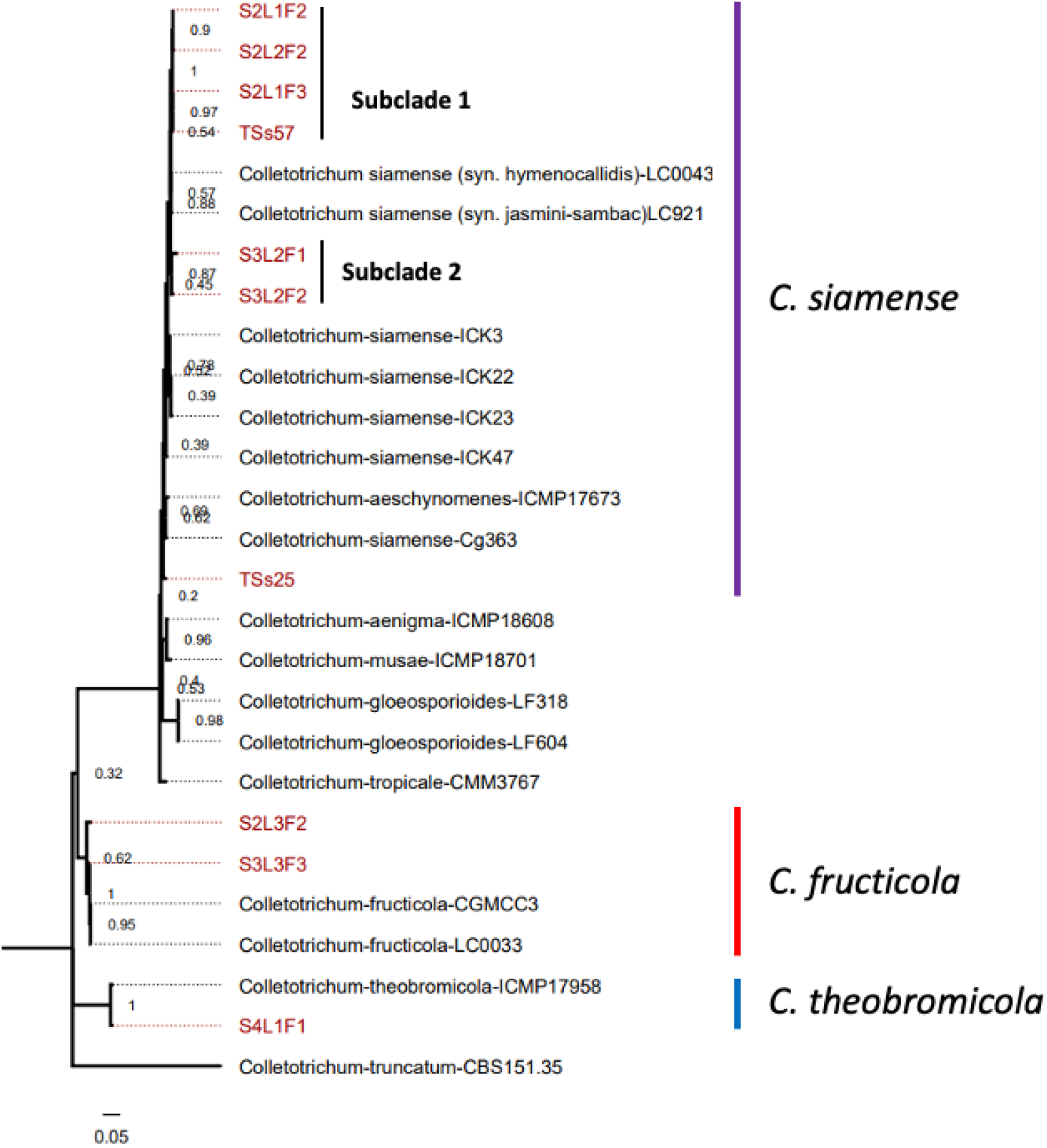
*Colletotrichum gloeosporioides* species complex phylogeny inferred by maximum parsimony from concatenated sequences of the ITS, ApMAT, GAPDH, and CAL loci. The 10 isolates from the current study are highlighted in red. Labels in black are for isolates for which sequences were retrieved from GenBank. The image represents the most parsimonious tree. The bootstraps support values are shown at the nodes. The tree is rooted with *C. truncatum* as the outgroup.

## Discussion

A disease survey was conducted to identify fungal diseases associated with African Yam Bean (AYB) in five states of Nigeria (Oyo, Enugu, Ebonyi, Abia, and Cross River; Fig. 1). A polyphasic approach was employed to identify the causative pathogens of the encountered diseases through morphological, molecular, and pathogenicity assays. During the field survey, leaf tip dieback, flower bud rot, and pod blight diseases were consistently observed (Fig. 2). Farmers who were interviewed during the survey testified to the adverse impact of these diseases on AYB production. Overall, this laboratory investigation determined that these diseases are caused by fungi belonging to four *Colletotrichum* species.

Fungal isolation revealed the presence of eight fungal genera associated with AYB diseases: *Botrydiplodia, Colletotrichum, Curvularia, Drechslera, Exserohilum, Fusarium*, *Pestalotia*, and *Seridium.* However, *Colletotrichum* spp., *Curvularia* spp., *Pestalotia* spp., and *Fusarium* spp., were more frequently isolated from the symptomatic tissues (Fig. 3). Afolabi et al. (2019) also identified *Colletotrichum*, *Curvularia*, *Fusarium*, and *Pestalotia*, as fungi associated with AYB flower bud rot and pod blight diseases. In that study, 20 AYB accessions were grown in a research field in Abeokuta, Nigeria during 2016/2017 and evaluated for their susceptibilities to flower bud and pod rot under natural conditions.

While *Curvularia* spp. and *Pestalotia* spp. are recognized as pathogens in Barbados lily and Tea plants (Liang et al. 2018; Liu et al. 2017), and *Fusarium* spp. are known as soilborne pathogens in cultivated plants (Arie 2019), our study found that the isolates of these fungi did not exhibit pathogenicity towards AYB. In contrast, nearly all isolates from the *Cg* species complex and all *C. truncatum* isolates tested in our laboratory assays induced disease symptoms. These findings implicate *Colletotrichum* spp. as significant causal agents of pod and leaf blights in AYB within Nigeria. The presence of *Curvularia* spp., *Pestalotia* spp., and *Fusarium* spp. in our samples raises questions about their roles in the AYB environment. They may exist as endophytes, potential sources of secondary infections, or saprophytes. To fully ascertain the pathogenicity of these fungi towards AYB, future studies should aim to inoculate AYB grown under field conditions with *Curvularia*, *Pestalotia*, and *Fusarium* isolates using diverse inoculation methods. These methods may include soil drenching, foliar spraying, or seed inoculation. It is crucial to note that detached leaf and pod assays, while informative, may not fully capture the complexity of interactions between these pathogens and AYB. Therefore, further investigation under field conditions or controlled environment (e.g. greenhouse) using diverse inoculation methods is necessary to thoroughly understand the risk posed by these fungi to AYB crops.

In the current study, we found that only *Colletotrichum* spp. were pathogenic. It is unclear if fungi not found in the current study but detected by Afolabi et al. (2019) (i.e., *Aspergillus*, *Cercospora*, *Macrophomina*, *Mucor*, *Pestalotia*, *Penicillium*, *Phomopsis*, *Pythium,* and *Rhizopus*) are also pathogenic to AYB. Additional research efforts are needed to improve the understanding of AYB pathogen diversity in Nigeria and elsewhere.

*Colletotrichum* is a genus composed of destructive pathogens responsible for immense losses in crop production. They cause diseases in leaves, stems, tubers, and fruits (Canon et al. 2012; Dean et al. 2012). Different complexes within the *Colletotrichum* genus exist, such as the *Cg* species complex (Cai et al. 2011; Weir et al. 2012). Employing morphological characters alone to delineate *Colletotrichum* spp. is inadequate as it does not provide enough information to differentiate among them effectively. For instance, in the current study, variations in colony morphology on PDA were observed within a species (*C. siamense* S2L1F3 and S2L1F2). Also, conidial size and radial growth measurement were similar among the isolates from the *Cg* species complex (Suppl. Fig. 1), although conidial shape, size and the presence of setae were useful in discriminating between *C. truncatum* and the *Cg* species complex (Suppl. Table 4). The appressorium of examined isolates was not informative in differentiating the species, as reported by Cai et al. (2009) and Sanders and Korsten (2003).

*C. truncatum* was observed to be a relatively weak pathogen on leaves compared to the *Cg* species complex (Fig. 6; Suppl. Table 3). Given high disease symptoms in pod pathogenicity assays and prevalence during pod isolations, the results suggest that *C. truncatum* from the current study are specialized to infect AYB pods (Fig. 5). Previous studies have shown that tissue specific infection patterns are common in fungal pathogens and may be caused by their adaptation to plant organs, available nutrients (Abrahamian et al. 2016), the developmental stages of the host (Barrett and Heil 2012) and pathogen, or the plant defense response (Lacaze and Joly 2020). Reasons for specialization of *C. truncatum* to preferentially infect AYB pods requires further investigation.

The ITS phylogenetic analysis allowed resolution of isolates into *Cg* species complex and *C. truncatum* but was inadequate to discriminate isolates within the *Cg* species complex. Similar observations were reported by Cai et al. (2009). The ITS, CAL, and GAPDH phylogenetic analysis, on the other hand, revealed that the isolates responsible for the observed AYB diseases belong to four different *Colletotrichum* species including *C. siamense* (comprising the largest number of isolates), *C. theobromicola*, *C. fructicola,* and *C. truncatum*. The three loci analysis was useful in separating *C. truncatum* and *C. theobromicola* (Fig. 9). The four loci phylogenetic analysis, however, provided a finer resolution, as it was effective in differentiating *C. fructicola* from the *C. siamense sensu lato* group (Fig. 10). It also provided branches with over all good boot-strap support values. The phylogenetic analysis revealed that in the *C. siamense* clade, isolates from Cross River (S2L1F2, S2L1F3, and S2L2F2) formed a distinct subclade (subclade 1) while isolates from Ebonyi (S3L2F1 and S3L2F2) clustered in subclade 2 (Fig. 10). Interestingly, isolate TSs57 from Oyo clustered with isolates in subclade 1 despite originating from a relatively far away distance. Long distance dispersal of a plant pathogen can occur through various mechanisms, including human-mediated transportation, air, large water bodies, and plant transmission (Golan and Pringle 2017; Nathan 2001).

There was no consistent trend on disease scores among the evaluated isolates of *C. theobromicola*, *C. fructicola*, and *C. siamense* (Fig. 4). Further, the two *C. siamense* isolates forming subclade 2 (Fig. 10), S3L2F1 and S3L2F2, had contrasting abilities to cause disease in the unwounded leaves used in the DLA. S3L2F1 did not cause disease while S3L2F2 had the highest score (Fig. 4). On the other hand, S3L2F1 had a high disease score when inoculated in wounded leaves (data not shown). Isolates of some *Colletotrichum* species are found to be pathogenic only when a host is wounded (Pring et al. 1995, Than et al. 2008). Testing a larger collection of *Cg* species complex isolates in their abilities to cause leaf blight in a diverse set of AYB accessions, using both wounded and unwounded leaves, warrants investigation to understand pathogen variability and mechanisms of resistance among AYB accessions.

The current study revealed the presence of four *Colletotrichum* species as causative pathogens of AYB foliar and pod diseases. These species have been characterized and reported on several hosts, highlighting their adaptability and potential impact across different cropping systems. For instance, *C. siamense*, *C. theobromicola*, and *C. fructicola* can cause anthracnose on coffee and mango, among other crops (Sharma et al. 2013; Weir et al. 2012) while *C. truncatum* causes anthracnose on soybean, pepper, papaya, among other crops (Boufleur et al. 2021; Torres-Calzada et al. 2018). To the best of our knowledge, this study is the most thorough examination of *Colletotrichum* species that impact AYB. Notably, it is the first documentation of *C. siamense*, *C. theobromicola*, and *C. fructicola* as the causative agents of AYB leaf diseases, as well as the first identification of *C. truncatum* as the pathogen that causes pod blight disease in AYB. To accurately delineate *Cg* species complexes, multi-locus sequencing has been the standard approach used. However, this method is time-consuming and requires refinement. Results from previous and the current study suggest that the ApMAT locus is a reliable marker for delimiting *Cg* species complex compared to other markers. Therefore, we suggest screening the ApMAT gene for classifying isolates within the *Cg* species complex.

Accurate identification of diseases is the first crucial step in designing disease management strategies. The presence of several *Colletotrichum* species associated with AYB diseases investigated in the current study may hinder decisions relating to disease management, with species possibly differing in their response to different management strategies or crop-resistance. Thus, further understanding of the plant-pathogen interactions involved in infection is critical to understand host resistance. Furthermore, understanding the response of these taxa to fungicides and alternative treatments will support development of integrated management strategies to control the diseases that they cause. Ultimately this will allow greater cultivation of this underutilized security legume crop which has substantial nutritional and nutraceutical promise.

## ACKNOWLEDGMENTS

We would like to express our sincere gratitude to the farmers and students (Mr. Olomitutu Oluwaseyi and Mr. Jeffrey Iheanacho) that provided access to their farms/research fields and allowed us to collect diseased tissues for conducting this research. The technical staff in the Pathology and Bioscience Units at IITA, Genetic Resources Centre, Faculty of Engineering and Science (UK), and Natural Resources Institute (UK) are appreciated for their support, particularly Mr. Greg Ogbe, Mr. Olalekan Ayinde, and Dr. Billy Ferrara. Olaide Ogunsanya expresses her gratitude to Dr. Yvonne Becker and Dr. Wolfgang Maier for their contributions in refining the scholarship proposal.

## AUTHOR’S CONTRIBUTION

Olaide Ogunsanya: Analysis (statistical and bioinformatics), investigation, methodology, writing original draft, writing review, and editing.

Clement Afolabi: conceptualization, methodology, supervision, writing-review and editing. Moruf Adebisi: supervision, writing-review and editing.

Akinola Popoola: supervision, writing-review and editing.

Olaniyi Oyatomi: methodology, supervision, writing-review and editing.

Richard Colgan: writing-review and editing.

Andrew Armitage: bioinformatics analysis, methodology, supervision, resources, writing-review and editing.

Elinor Thompson: methodology, supervision, resources, writing-review and editing.

Michael Abberton: Funding acquisition, project administration, resources, supervision, writing-review.

Alejandro Ortega-Beltran: methodology, supervision, resources, writing-review, and editing.

