## Supplementary material for "Morphological, pathological and phylogenetic analyses identify a diverse group of *Colletotrichum* spp. causing leaf, pod, and flower diseases on the orphan legume African yam bean": Suppl. Table 1

**Supplementary Table 1. Origin, host crop, and GenBank accession numbers of sequences of four loci of Colletotrichum isolates used in the current study.**

| Fungus | Isolate code | Origin | Host crop | GenBank accession number | | | |
| --- | --- | --- | --- | --- | --- | --- | --- |
|  |  |  |  | ITS^a^ | GAPDH^a^ | CAL^a^ | APMAT^a^ |
| *C. siamense* | ICK-3 | Korea | Persimmon | LC260488 | LC260529 | LC260532 | LC307171 |
|  | ICK-22 | Korea | Persimmon | LC260489 | LC260530 | LC260533 | LC307172 |
|  | ICK-23 | Korea | Persimmon | LC260490 | LC260531 | LC260534 | LC307173 |
|  | ICK-47 | Korea | Persimmon | LC208833 | LC208834 | LC208835 | LC307174 |
|  | Cg363 | Japan | Strawberry | NW-023501017 | NW-023501015 | NW-023501018 | NW-023501021 |
| *C. hymenocallidis* | ICMP 18642 | China | *Hymenocallis littoralis* | JX010278 | JX010019 | JX009709 | JQ899283 |
| *C. jasmini-sambac* | LC921 | Vietnam | *Jasminum sambac* | HM131511 | HM131497 | HM131492 | JQ807841 |
|  | S3L2F1 | Nigeria | Africa yam bean | OQ586196 | OQ581982 | OQ581977 | OQ581963 |
|  | S2L1F2 | Nigeria | Africa yam bean | OQ586192 | OQ581988 | OQ581971 | OQ581959 |
|  | S2L1F3 | Nigeria | Africa yam bean | OQ586193 | OQ581992 | OQ581967 | OQ581960 |
|  | TSs 57 | Nigeria | Africa yam bean | OQ586189 | OQ581994 | OQ581974 | OQ581956 |
|  | TSs 25 | Nigeria | Africa yam bean | OQ586190 | OQ581995 | OQ581973 | OQ581957 |
|  | S3L1F1 | Nigeria | Africa yam bean | OQ586197 | OQ581989 | OQ581975 | - |
|  | S2L2F2 | Nigeria | Africa yam bean | OQ586188 | OQ581986 | OQ581968 | OQ581955 |
|  | S3L2F2 | Nigeria | Africa yam bean | OQ586195 | OQ581980 | OQ581965 | OQ581962 |
| *C. fructicola* | CGMCC3 | China | Strawberry | NW-0224744931 | NW-022474561 | NW-022474237 | NW-022474248 |
|  | ICMP 18581 | Thailand | *Coffea arabica* | JX010165 | JX010033 | FJ917508 | JQ807838 |
|  | S2L3F2 | Nigeria | Africa yam bean | OQ586194 | OQ581979 | OQ581966 | OQ581961 |
|  | S3L3F3 | Nigeria | Africa yam bean | OQ586191 | OQ581987 | OQ581972 | OQ581958 |
| *C. aenigma* | CG56 | Japan | Strawberry | NW-023500921 | NW-023500908 | NW-023500911 | NW-023500914 |
|  | ICMP 18608 | Israel | *Persea american* | JX010244 | JX010044 | JX009683 | KM360143 |
| *C. nupharicola* | CBS 470 | USA | *Nuphar lutea* | JX145173 | JX009972 | JX009663 | JX145319 |
|  | CBS 469.96; ICMP 17938 | USA | *Nuphar lutea subsp. polysepala* | JX010189 | JX009936 | JX009661 | - |
| *C. gloeosporioides* | MTCC 10323 | India | *Citruslinensis* | KC790935 | - | - | JQ807843 |
|  | LC1 | China | *Liriodendro chinensis* | NW-025544752 | NW-025544729 | NW-025544669 | NW-025544710 |
| *C. theobromicola* | ICMP 17958 | Australia | *Stylosanthes guianensis* | JX010291 | JX009948 | JX009598 | - |
|  | S4L1F1 | Nigeria | Africa yam bean | OQ586187 | OQ581990 | OQ581976 | OQ581954 |
| *C. truncatum* | CBS 151.35 | USA | *Phaseolus lunatus* | MH855611 | GU228254 | KY856132 | **-** |
|  | TSs 432 | Nigeria | Africa yam bean | OQ586200 | OQ581985 | OQ581978 | **-** |
|  | Pod 6 | Nigeria | Africa yam bean | OQ586198 | OQ581991 | OQ581969 | **-** |
|  | TSs 29 | Nigeria | Africa yam bean | OQ586199 | OQ581984 | OQ581970 | **-** |
| *C. higginsianum* | C5 | China | *Rumex acetosa* | MF033888 | MF033889 | MF033893 | **-** |
| *C. tropicale* | CMM3767 | Brazil | *Magnifera indica* | KC702985 | KC702960 | KC992378 | KJ155464 |
| *C. aeschynomenes* | ICMP 17673 | USA | *Aeschynomene virginca* | JX010176 | JX009930 | JX009721 | KM360145 |
| *C. musae* | ICMP 18701 | Philippines | *Musa* spp. | JX010145 | JX010047 | JX009687 | - |

^a^ ITS: internal transcribed spacer; GAPDH: glyceraldehyde 3- phosphate dehydrogenase; CAL: calmodulin; ApMAT: Apn2-MAT 1-2 intergenic spacer
