## Supplementary material for "Morphological, pathological and phylogenetic analyses identify a diverse group of *Colletotrichum* spp. causing leaf, pod, and flower diseases on the orphan legume African yam bean": Suppl. Table 2

**Supplementary Table 2. Origin of *Colletotrichum gloeosporioides* species complex evaluated in their abilities to cause foliar disease of African yam bean in detached leaf assays.**

| Isolate | Origin | Local Government Area | Geopolitical zone | Latitude (North) | Longitude (East) | Elevation (m) |
| --- | --- | --- | --- | --- | --- | --- |
| S2L1F2 | Cross River | Ogoja | South South | 6.64973 | 8.78504 | 64 |
| S2L1F3 | Cross River | Ogoja | South South | 6.58341 | 8.79834 | 96 |
| S2L2F2 | Cross River | Yala | South South | 6.66268 | 8.62770 | 57 |
| S2L3F2 | Cross River | Bekwarra | South South | 6.67852 | 8.96469 | 129 |
| S3L1F1 | Ebonyi | Afikpo North | South East | 5.95185 | 7.95664 | 161 |
| S3L1F2 | Ebonyi | Afikpo North | South East | 5.95052 | 7.95377 | 36 |
| S3L2F1 | Ebonyi | Afikpo South | South East | 5.87870 | 7.77881 | 41 |
| S3L2F2 | Ebonyi | Afikpo South | South East | 5.87862 | 7.77892 | 39 |
| S3L3F1 | Ebonyi | Onicha | South East | 6.02473 | 7.91977 | 34 |
| S3L3F3 | Ebonyi | Onicha | South East | 6.00173 | 8.00021 | 20 |
| S4L1F1 | Enugu | Nkanu East | South East | 6.26101 | 7.60909 | 121 |
| S4L1F3 | Enugu | Nkanu East | South East | 6.26201 | 7.61260 | 80 |
| S4L2F1 | Enugu | Enugu South | South East | 6.37478 | 7.49893 | 182 |
| S4L2F3 | Enugu | Enugu South | South East | 6.37262 | 7.53778 | 133 |
| TSs 25 | Oyo | Ibadan | South West | 7.49620 | 3.90760 | 230 |
