## Supplementary material for "Morphological, pathological and phylogenetic analyses identify a diverse group of *Colletotrichum* spp. causing leaf, pod, and flower diseases on the orphan legume African yam bean": Suppl. Table 3

**Supplementary Table 3. Comparison of disease score values among isolates belonging to *Colletotrichum truncatum* and *Cg* species complex when inoculated on African yam bean leaves.**

| **ISOLATES^a^** | **DS4DAI^a^** | **MR-DS4DAI ^b^** | **DS7DAI^b^** | **MR-DS7DAI ^c^** | **DS10DAI ^b^** | **MR-DS10DAI ^c^** |
| --- | --- | --- | --- | --- | --- | --- |
| CONTROL | 1 | 5 | 1 | 4.5 | 1 | 4 |
| **TSs 98** | 2 | 17 | 2 | 15.5 | 2 | 15 |
| **TSs 432c** | 1.3 | 9 | 1.3 | 8.2 | 1.3 | 7.7 |
| **Pod 6** | 1 | 5 | 1.3 | 8.2 | 1.3 | 7.7 |
| **TSs 29B** | 2 | 17 | 2 | 15.5 | 2 | 15 |
| **TSs 421** | 2 | 17 | 2.7 | 20.8 | 2.7 | 19.3 |
| **TSs 61** | 2 | 17 | 2 | 15.5 | 2 | 15 |
| S2L2F2 | 3 | 29 | 3.3 | 27.8 | 3.7 | 26.8 |
| S2L1F3 | 2.3 | 21 | 2.3 | 18.8 | 2.7 | 21.3 |
| S2L3F2 | 2.3 | 21 | 3 | 24.3 | 3.3 | 23.7 |
| S2L1F2 | 3 | 29 | 3.3 | 27.8 | 5 | 31.5 |
| ***P*-value** | ** |  | * |  | ** |  |
| **H^d^** | 24.89 |  | 22.33 |  | 26.26 |  |

^a^ The mean severity score of isolates within the *C. truncatum* and *Cg* species complex were evaluated for pathogenicity.

^b^ DS4DAI – DS8DAI= Disease score 4 - 8days after inoculation

^c^ MR-DS4DAI – DS8DAI= Mean rank of disease score 4 - 8days after inoculation

^d^ H is the Chi-Square value

*C. truncatum* isolates are in bold.
