## Supplementary material for "Morphological, pathological and phylogenetic analyses identify a diverse group of *Colletotrichum* spp. causing leaf, pod, and flower diseases on the orphan legume African yam bean": Suppl. Table 4

**Supplementary Table 4. Morphological features of Colletotrichum isolates belonging to various species that were collected from diseased African yam bean tissues.**

| **Criterion** | ***Colletotrichum jasmini-sambac* (S2L1F2)** | ***Colletotrichum fructicola* (S2L3F2)** | ***Colletotrichum theobromicola* (S4L1F1)** | ***Colletotrichum truncatum* (TSs 29)** |
| --- | --- | --- | --- | --- |
| Colour on PDA | Aerial mycelia are olivaceous with rings of orange and black conidia masses | Aerial mycelia are olivaceous grey with few orange conidia masses | Olivaceous aerial mycelia | whitish-grey aerial mycelia |
| Growth rate on PDA | Fast growing | Fast growing | Fast growing | Intermediate growing |
| Conidia shape | One-celled, aseptate, cylindrical with rounded ends | One-celled, aseptate, cylindrical with rounded ends | One-celled, aseptate, cylindrical with rounded ends | One-celled, aseptate, and falcate |
| Conidia size | 5.0 to 15.0 µm × 2.5 to 7.5 µm | 7.5 to 12.5 µm × 2.5 to 5.0 µm | 10.0 to 22.5 × 2.5 to 5.0 µm | 19.0 to 26.5 × 3.5–4.5 µm |
| Appressoria shape | Brown to dark brown with irregular margins | Brown to dark brown, sub-fusoid, lobed to irregular margins | Irregular, dark to brown | Irregular, dark to brown |
| Setae | Absent | Absent | Absent | Present |
