## Supplementary material for "Morphological, pathological and phylogenetic analyses identify a diverse group of *Colletotrichum* spp. causing leaf, pod, and flower diseases on the orphan legume African yam bean": Suppl. Fig. 1

Supplementary Figure 1. Conidia of *Colletotrichum* species isolated from diseased AYB tissues. A – D is showing conidia shape of isolates from the *C. gloeosporioides* species complex (A. S2L1F2, B. S2L1F3, C. S2L3F2, D. S4L1F1) while E – F is showing conidia shape of isolates from C*. truncatum* (E. TSs 432, F. TSs 29; Scale bar of A - F =20 μm). Magnification is shown at 40×.

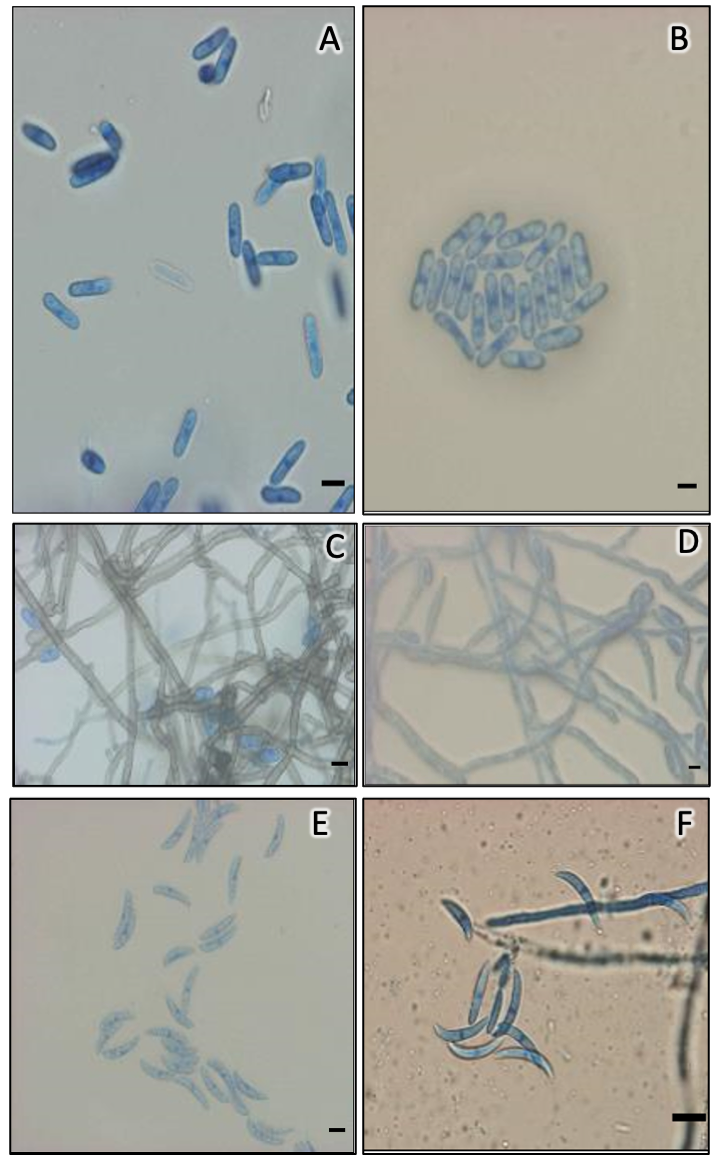
