## Supplementary material for "Morphological, pathological and phylogenetic analyses identify a diverse group of *Colletotrichum* spp. causing leaf, pod, and flower diseases on the orphan legume African yam bean": Suppl. Fig. 2

Supplementary Figure 2. Variation in conidia size of *Colletotrichum* spp. isolated from African yam bean leaves, and pods.

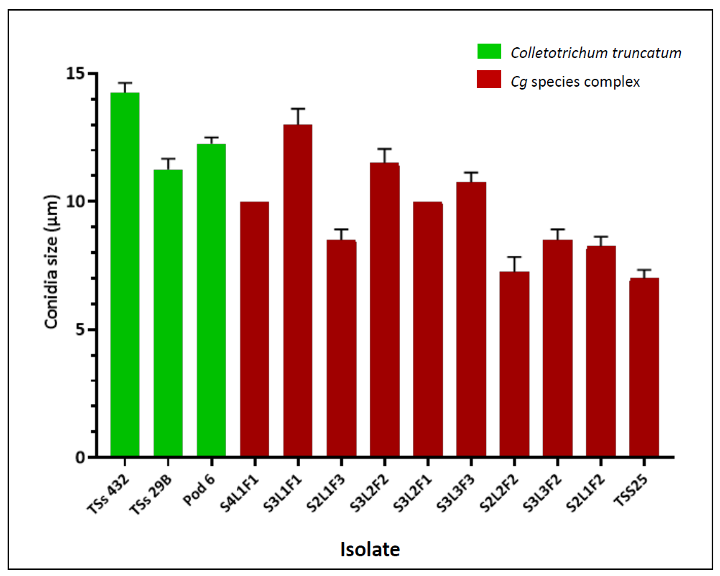
