## Supplementary material for "Morphological, pathological and phylogenetic analyses identify a diverse group of *Colletotrichum* spp. causing leaf, pod, and flower diseases on the orphan legume African yam bean": Suppl. Fig. 3

Supplementary Figure 3. Morphological features of *Colletotrichum* species isolated from diseased AYB tissues. A – C show appressorium shape of isolates from the *C. gloeosporioides* species complex (A. S2L1F3, B. S2L3F2, C. S4L1F1), D shows appressorium shape of *C. truncatum* isolate TSs432 while E – F is showing setae produced by C*. truncatum* isolates (E. TSs 432, F. TSs 29). Scale bar of A - F =20 μm at 40× magnification.

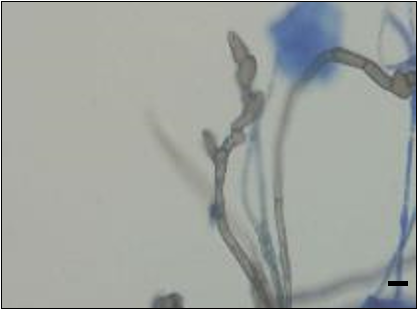

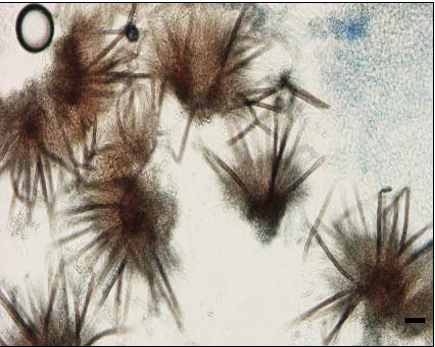

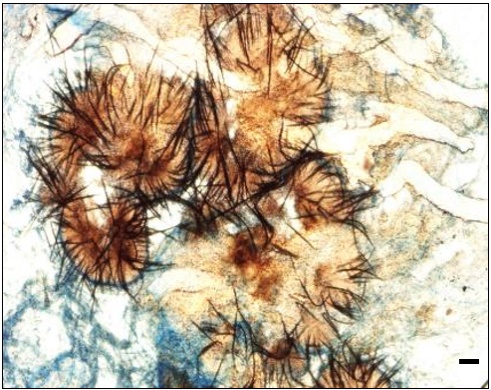

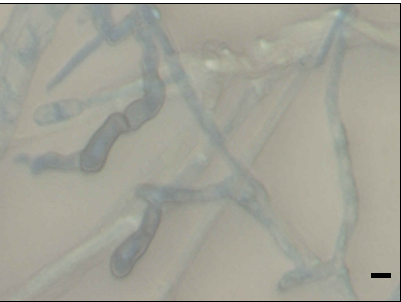

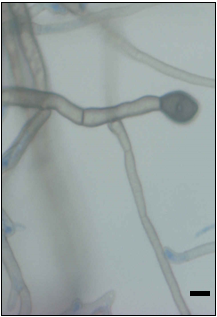

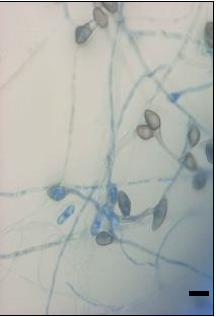

A

C

E

D

F

B
